# URAdime – a tool for analyzing primer sequences in sequencing data to identify dimers and super-amplicons

**DOI:** 10.1101/2025.01.25.634870

**Authors:** Jason D Limberis, John Z Metcalfe

**Affiliations:** Division of Pulmonary and Critical Care Medicine, Zuckerberg San Francisco General Hospital and Trauma Centre, University of California, San Francisco, San Francisco, CA, USA

## Abstract

Targeted sequencing of multiplex PCR amplicons is commonplace in research laboratories and clinical diagnostics. There are numerous tools for the a priori optimization of primers and reactions, but no tools to detect specific problematic primers post hoc. We developed URAdime, a tool for analyzing primer sequences in sequencing data to identify dimers and super-amplicons. We show that it successfully detects these unwanted amplicons and accurately attributes their generation to specific primers.

## Introduction

Targeted sequencing involves multiplex PCR for specific targets followed by sequencing on various platforms and is commonplace in molecular biology and clinical diagnostics. Multiplex PCR is the simultaneous amplification of numerous targets (≥ 2) within the same reaction; the reaction is influenced by numerous factors, including the primer sequences (GC content, specificity, annealing temperature, secondary structure formation, concentration), buffer components, and the polymerase used. Multiple tools exist(Untergasser *et al*., 2012) to design single primer pairs and detect primer specificity and secondary formation, and even some to rationally design multiplexed PCR primers(Limberis and Metcalfe, 2023; Kechin *et al*., 2020; Xie *et al*., 2022). While these tools are excellent and provide *a priori* optimization, they cannot model the entire reaction. Thus, primer design and reaction conditions often need to be augmented based on empirical results. There are also a few tools for chimeric amplicon detection(Wright *et al*., 2012; Edgar, 2016) like UCHIME2, which identify and filter out artifact sequences but are designed to filter out reads arising from the same primer set caused by a fusing between two different amplicons (e.g., generated when an incomplete amplicon from one species acts as a primer for amplicons for a different species causing a fusion). These artifacts can occur during PCR, especially during metagenomic sequencing. No published tools specifically quantify the primer-dimers and super-amplicons in a targeted sequencing reaction. Super-amplicons occur when primer pairs are skipped during amplification, resulting in longer-than-expected products when distant primers incorrectly pair. We present URAdime (Universal Read Analysis of DIMErs), a simple tool for analyzing primers in sequencing data to identify primer-dimers and super-amplicons. URAdime allows researchers to identify problematic primers to aid iterative optimization of the primer sets and reaction conditions.

## Methods and Usage

URAdime is written in Python and operates as both a command-line tool and as a Python package. URAdime accepts a reference sequence file and a bam file; it is agnostic to the sequencing technology used (however, the full amplicon size cannot be reliably determined from short-read data). Primer information is supplied in a tab-separated format specifying primer pair names, forward and reverse sequences, and expected amplicon sizes (see README). The implementation utilizes libraries including pysam for BAM file processing, BioPython for sequence manipulation, and python-Levenshtein for efficient sequence similarity calculations.

URAdime converts the input bam file to fasta sequences [--downsample percentage, range: 0.1-100.0%] and then searches for the input primers in the 5’-ends of the sequence [--window-size, default: 20 bp] through a Levenshtein distance-based matching algorithm [--max-distance Levenshtein distance, default: 2 and overlap requirements --overlap-threshold, default: 0.8] (**Figure 1**). The window-size is also useful when multiplex primer sets contain universal tail sequences that would shift the reads primer position. URAdime can also be restricted to analyzing only unaligned reads [--unaligned-only], as these are likely unexpected amplicons depending on the initial alignment parameters. URAdime implements a flexible size tolerance system to validate amplicon sizes [size-tolerance, default: 10%] with a bypass option [--ignore-amplicon-size]. The tool can also output a filtered BAM file [--filtered-bam] containing only correctly matched and sized reads for downstream analysis. The analysis implements multi-threading capabilities [-t threads, default: 4] with configurable chunk sizes for parallel processing [-c, default: 100 reads per chunk].

**Figure 1:**
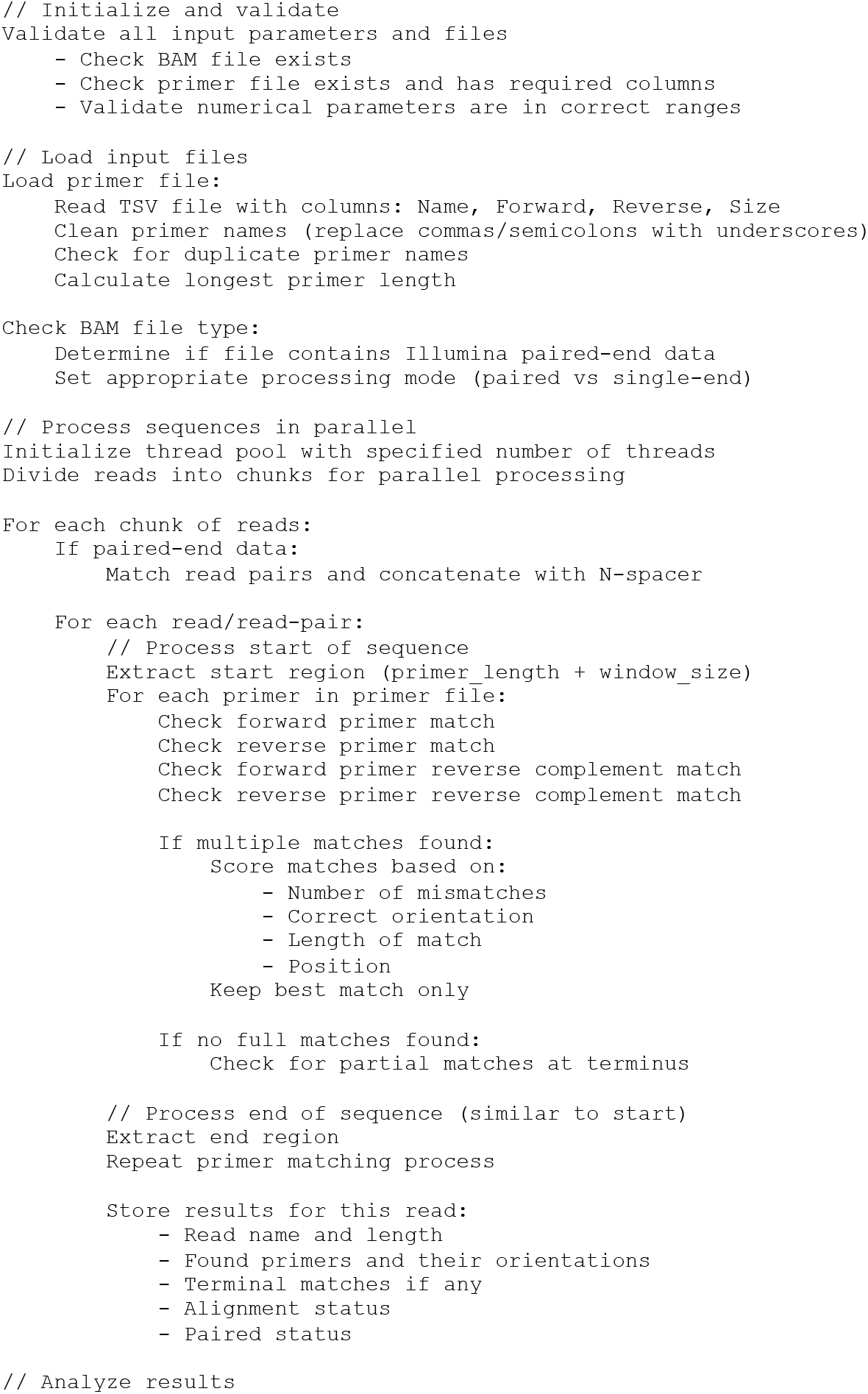
URAdime pseudocode, see codebase for detailed functions and descriptions.

URAdime outputs a summary counting the number of sequences that have no primers, have only one primer, have correct matching primer pairs (correct primers at both ends, in the proper orientation, with sequence length matching the expected size [only possible for long-read sequencing]), and have wrong matching primer pairs (wrong primers or wrong size). It also outputs files containing the read names and details about the primers found, including their location and orientation. Finally, URAdime outputs per primer summaries, allowing the user to identify problematic primers quickly. We also built a user-friendly web-based application to run URAdime (https://drdx.ucsf.edu/URAdime.html).

## Results

We created a helper script (generate_toy_dataset.py) to generate a synthetic test datasets. Primer sequences are generated using create_random_primer(), which constructs oligonucleotides of 18-25bp with controlled GC content (40-60%) using a probabilistic base selection approach.

The create_dataset_params() function defines the number of primer pairs (5-10), size ranges (100-1000bp), total read count (800-1200), and the category distributions. generate_read() generates the insert sequences with controlled GC content using a weighted random sampling approach. Reads are compiled into BAM format using pysam, with read tags corresponding to their expected. The generator creates twelve distinct read categories corresponding to those outputted by URAdime. URAdime successfully detected all categories with high accuracy (**Table 1**).

**Table 1:**
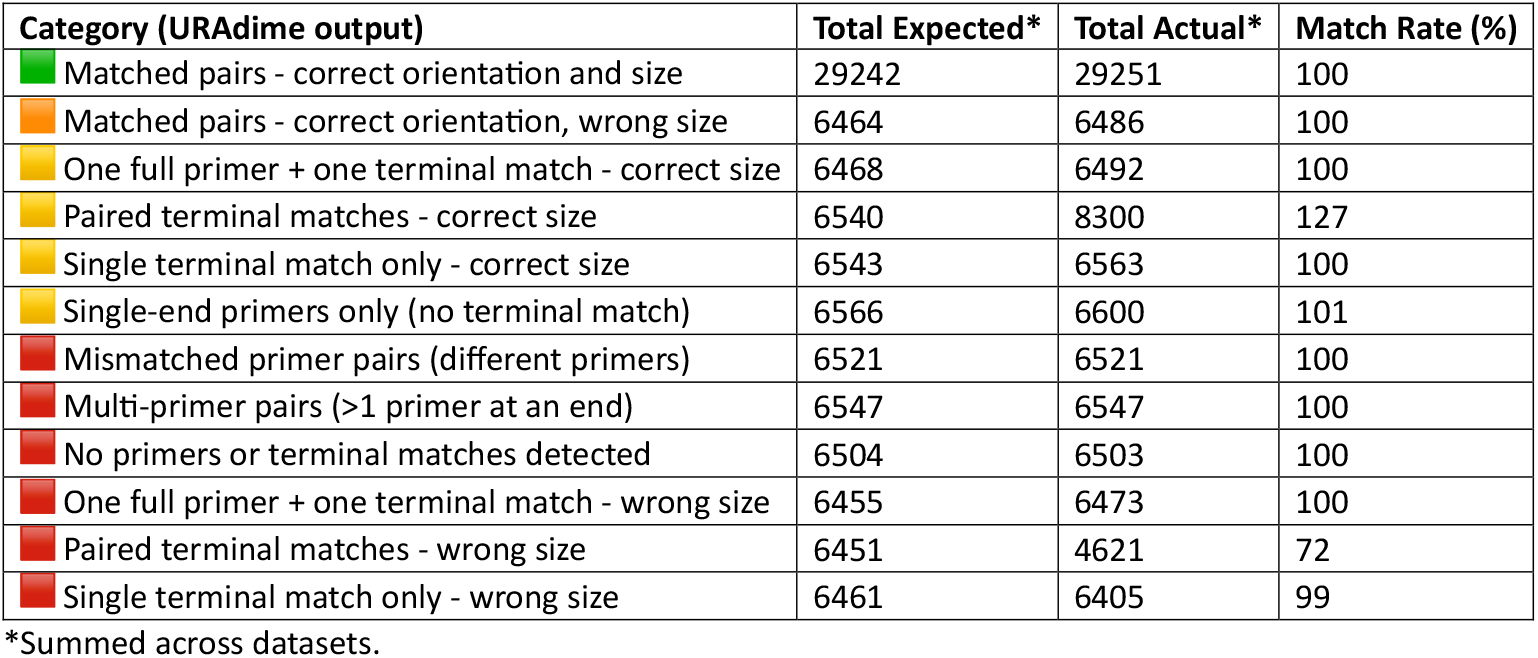
URAdime correctly detects the various amplicon species in a bam file generated using “generate_comprehensive_dataset.py” and run with edit distance set to 0 (--max-distance 0). The analysis includes 100 simulated datasets. The number of primer sets averaged 7 (SD 1.5; range 5 to 10) with a primer length averaging 22 (SD 1; range 20 to 22 nucleotides [the range within datasets was 18 to 25]). The primer GC content averaged 0.50 (SD 0.03; range 0.41 to 0.62). There were an average of 1008 (SD 113; range 800 to 1193) reads per sample. The minimum size of the amplicons averaged 155bp (SD 29; range 100 to 200bp) while the maximum size averaged 662 bp (SD 189; 307 to 992 bp). The amplicon GC content averaged 0.51 (SD 0.03; range 0.41 to 0.60). Full details are in the test folder (uradime_detailed_summary.xlxs).

| Category (URAdime output) | Total Expected* | Total Actual* | Match Rate (%) |
| --- | --- | --- | --- |
| Matched pairs - correct orientation and size | 29242 | 29251 | 100 |
| Matched pairs - correct orientation, wrong size | 6464 | 6486 | 100 |
| One full primer + one terminal match - correct size | 6468 | 6492 | 100 |
| Paired terminal matches - correct size | 6540 | 8300 | 127 |
| Single terminal match only - correct size | 6543 | 6563 | 100 |
| Single-end primers only (no terminal match) | 6566 | 6600 | 101 |
| Mismatched primer pairs (different primers) | 6521 | 6521 | 100 |
| Multi-primer pairs (>1 primer at an end) | 6547 | 6547 | 100 |
| No primers or terminal matches detected | 6504 | 6503 | 100 |
| One full primer + one terminal match - wrong size | 6455 | 6473 | 100 |
| Paired terminal matches - wrong size | 6451 | 4621 | 72 |
| Single terminal match only - wrong size | 6461 | 6405 | 99 |

\*Summed across datasets.

## Discussion

URAdime is a novel tool for analyzing sequencing data and identifying problematic primers within multiplex PCR experiments. The optimization of multiplex PCR is essential for accurate and efficient sequencing and is utilized in most research labs and many clinical diagnostic assays. This posthoc analysis complements a priori optimization tools, providing insights into specific primers’ performance under particular reaction conditions in real-world experiments. URAdime provides a user-friendly, efficient, and accurate method to detect unwanted amplicons enabling researchers to refine their primer design and can improve the success and efficiency of multiplex PCR.

## Data availability

URAdime source code and raw data are available at https://github.com/SemiQuant/URAdime, the URAdime package is available at https://pypi.org/project/URAdime/ and a server version is available for use at www.DrDx.Me.

